# Current Research On Epigenetic Age and Cellular Senescence: A Bibliometric and Visual Analysis

**DOI:** 10.1101/2024.02.13.580124

**Authors:** M.M Mureti kahaer, B.M Wang Hao, M.M Zhi-Guo Sun, M.M Kuo Xu, M.M Dilimulati Aikeremu

**Affiliations:** Second Ward of Spine, Orthopedic Center, Xinjiang Uygur Autonomous Region People’s Hospital, 830000, Urumqi, Xinjiang, Republic of China

**Keywords:** epigenetic age, bibliometrics, visualization analysis, Cell aging

## Abstract

The aim of this study is to summarize and visualize the current state of research on epigenetic age and cellular senescence over the last two decades using bibliometric methods and to explore the current research priorities and future development directions. Relevant literature on the topic Epigenetic age and cellular senescence data published between January 2000 and January 2024 were searched in the Web of Science core database and three bibliometric mapping tools (Biblimetrix R Package, VOS Viewer and CiteSpace) were used to provide an overview of the Literature to give and analyze keywords, the co-occurrence of reference citations, authors and institutions, to analyze the current status of epigenetic aging research and hot changes in recent years. A total of 2,193 relevant studies were included, and the co-occurrence network identified seven clusters of associations between epigenetic age and cellular senescence, including effects on human tissues, genetic clock analyses, the relationship of age-related diseases, precise genetic changes and accelerated cellular senescence, localized cellular transformations, and genetic immune regulation. The analysis shows that research in this area has shifted from disease-level studies to mechanism-based studies, with cancer being the main disease and the main pathogenic mechanism being related to DNA methylation. The research direction in recent years has focused on the mechanism study of epigenetic age and cellular senescence as well as target therapy. The bibliometric and visualization analyzes provide a comprehensive understanding of the research progress and the role between epigenetic age and human cellular senescence and enrich the review literature in this field.

## Introduction

Human cell senescence is a change process in which the cell’s ability to proliferate and differentiate, as well as its physiological functions, gradually decrease over time as the cell carries out its life activities [1]. Epigenetic clocks are age calculators derived using mathematical formulas [2] that use to estimate specific CpG methylation levels in the genome that change with age. The estimated age value is called epigenetic age. The actual age is called epigenetic acceleration [3]. A number of studies have linked annual epigenetic acceleration to a variety of chronic human diseases [4], human cellular health [5], lifestyle [6], mental state [7] and environmental factors [8] and have a connection bet ween epigenetic age and aging [9]. Although such findings have provided some theoretical insights into the causes of aging, the mechanisms by which epigeneticage regulates cellular senescence remain largely unclear [10]. The aim of this article is therefore to use bibliometrics and visualization analysis to review the published literature secondary to analyze, investigate the development trend of the relationship between epigenetic age and human cellular aging, and provide theoretical guidelines for clinical related researchers.

## Materials and methods

### 1.1 DATA COLLECTION

Web of Science is considered the most comprehensive and reliable database for bibliometric analysis, covering virtually all high-quality and influential journals and providing a comprehensive citation dataset. To improve data representativeness and accessibility, all data were extracted from the Web of Science Core Collection (https://www.webofscience.com/wos/woscc/). The search period was set from January 2000 to January 2024 using “epigenetic age” and “aging in human cells” “senescence” as catchphrases. In addition, the literature records were defined as complete records and cited references and fin ally exported to plain text format. Exclusion criteria: (1) non-English literature; (2) Conference abstracts, editorials, letters, book chapters, and other types of literature.

### 1.2 Bibliometrics analysis and visualization

#### 1.2.1 Using the Bibliometrix package in R software

the TXT data downloaded from WOS was imported to visualize and analyze the literature authors, citation frequency and country/region, institution, journal, etc.; The Impact Factors (IFs) used for journals were taken from the Journal Citation Reports (JCR) 2023.

#### 1.2.2 Co-citation analysis of the literature was performed using Cite Space co-occurrence analysis software

In all generated graphs, each node represents an element, including author, country, organization and keyword, and the frequencies are all represented by the size of the points. The lines connecting the dots rep resent relationships of co-occurrence or co-citation, while the number of connected lines represents the closeness of collaboration between the dots as well as the strength of co-occurrence or co-citation. The Cites pace software parameters were set as follows: selection of period (2000-2024), number of years in each period (2), terminology sources (select all), selection criteria (top 50%), visualization (staticcluster view, display merged networks).

#### 1.2.3 Cluster analysis of keywords was performed using VosViewer visualization software

Data were imported into VosViewer software, key terms were extracted from titles and abstracts, keywords with at least five occurrences were retained, and co-occurrence results were obtained by manually removing irrelevant keywords. In the visualization results, the elements involved in the keywords are represented as nodes in space defined, and the connecting lines between the nodes indicate the coreferential relationships between these elements. The related items are classified differently and the importance is indicated by different colors, shapes and sizes. In a co-citation graph, different points represent different keywords, and the size of the points corresponds to the number of elements. Lines connecting dots to dots indicate co-citation relationships. Different colored dots and lines indicate different clusters or time periods.

#### 1.2.4 All data and information in this article

were obtained from the Web of Science (WOS) open access literature database, so this study did not require informed consent and ethical approval from relevant institutions.

## Results

### 2.1 Global research status and trends of the epigenetic age

#### 2.1.1 Current volume of published literature worldwide

A total of 2139 English-language studies on epigenetic age in relation to human cell senescence were retrieved through the WOS core database. The annual volume of publications is shown in Fig 1. The result is Fig 1 shows that the number of publications on genetic age increased year by year from 2000 to 2024, with the highest number of publications in 2022 with 213 publications, accounting for 9.95% of the total number of publications (hence the number of publications in 2024). was not included in the comparison because the current study was completed in January 2024. The number of publications in 2024 was not included in the comparison.

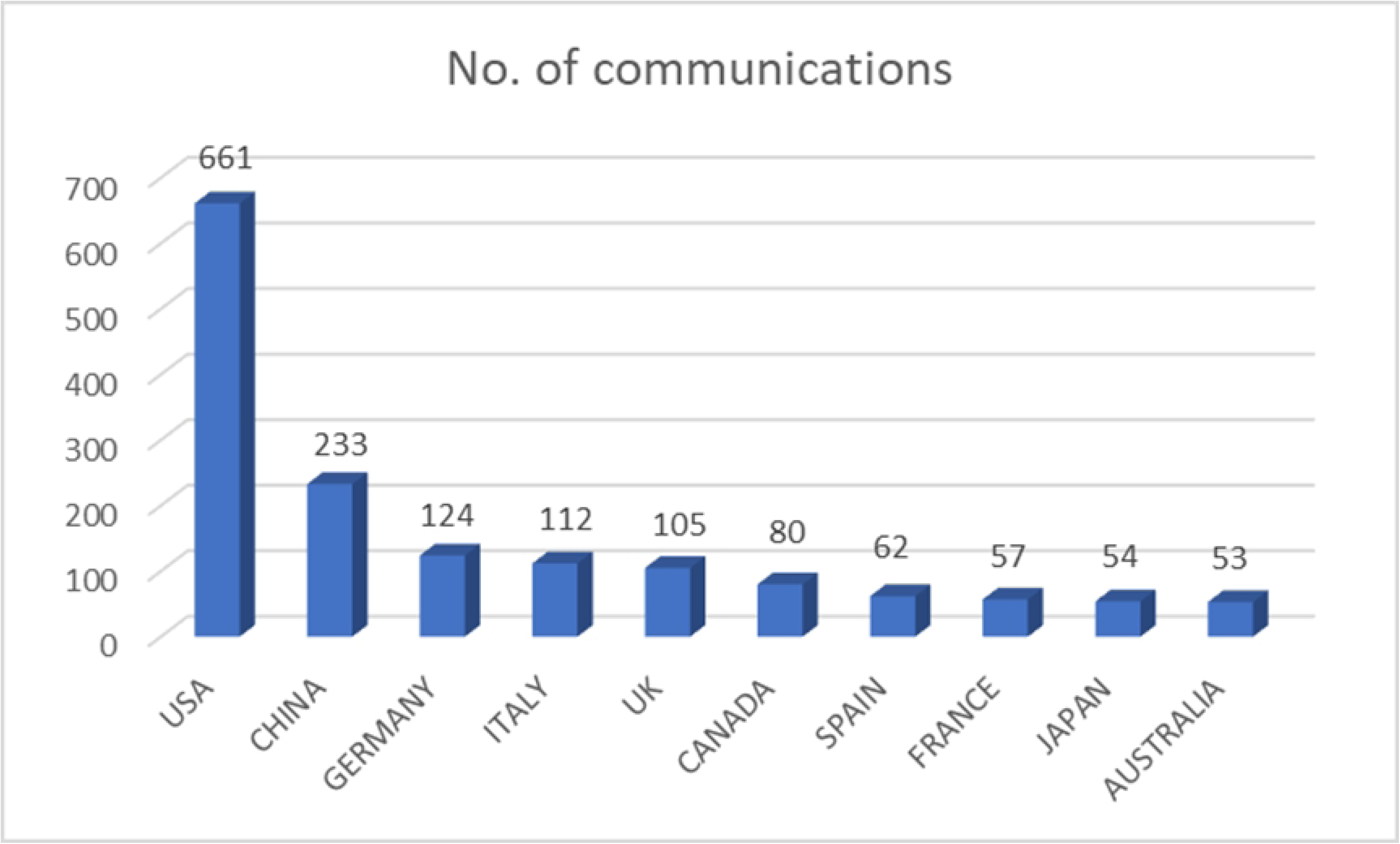

#### 2.1.2 Number of publications in different countries/regions

Visualization of collaboration between countries/regions on the online platform bibliometrix. Figs 2A and B shows that a total of 84 countries or regions have published studies on epigenetic age and human cellular aging during the period 2000–2024. and the statistics show that the United States took first place in the number of publications, with a total of 661 publications; China ranked second after the United States with 233 publications; Germany came third with 124 publications; Italy ranked fourth and fifth with 112 publications and the United Kingdom ranked fourth and fifth with 105 publications. The statistics show that the United States ranked first in terms of the number of publications, with a total of 661 publications; With a total of 233 publications, China took second place after the USA; Germany came third with 124 publications; Italy ranked fourth with 112 publications; and the United Kingdom came fourth and fifth with 105 releases.

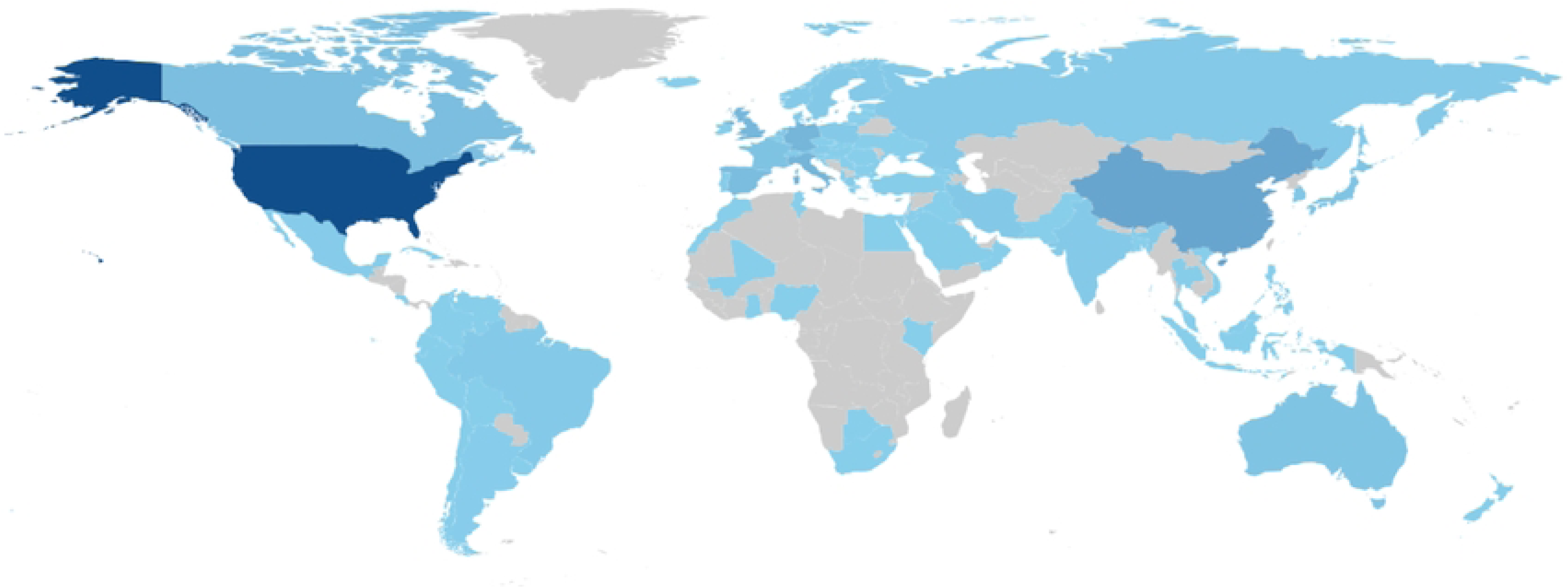

#### 2.1.3 Publication quality in different countries/regions

In bibliometrics, the total number of citations and the H-index are important data for assessing the quality of literature. In bibliometrics, the total number of citations and the H-index are important data for evaluating the quality of literature. The higher the total number of citations, the more the literature is accepted by more scholars and the higher the quality of the literature. (Fig 3) shows the top 5 countries/reg ions in epigenetic years, with the United States having the highest total number of citations (46,031), Spain (13,660) in second place, and Germany (6,837), China (6,188), and Canada (5,698) ranked third, fourth and fifth.

**Fig 3.**
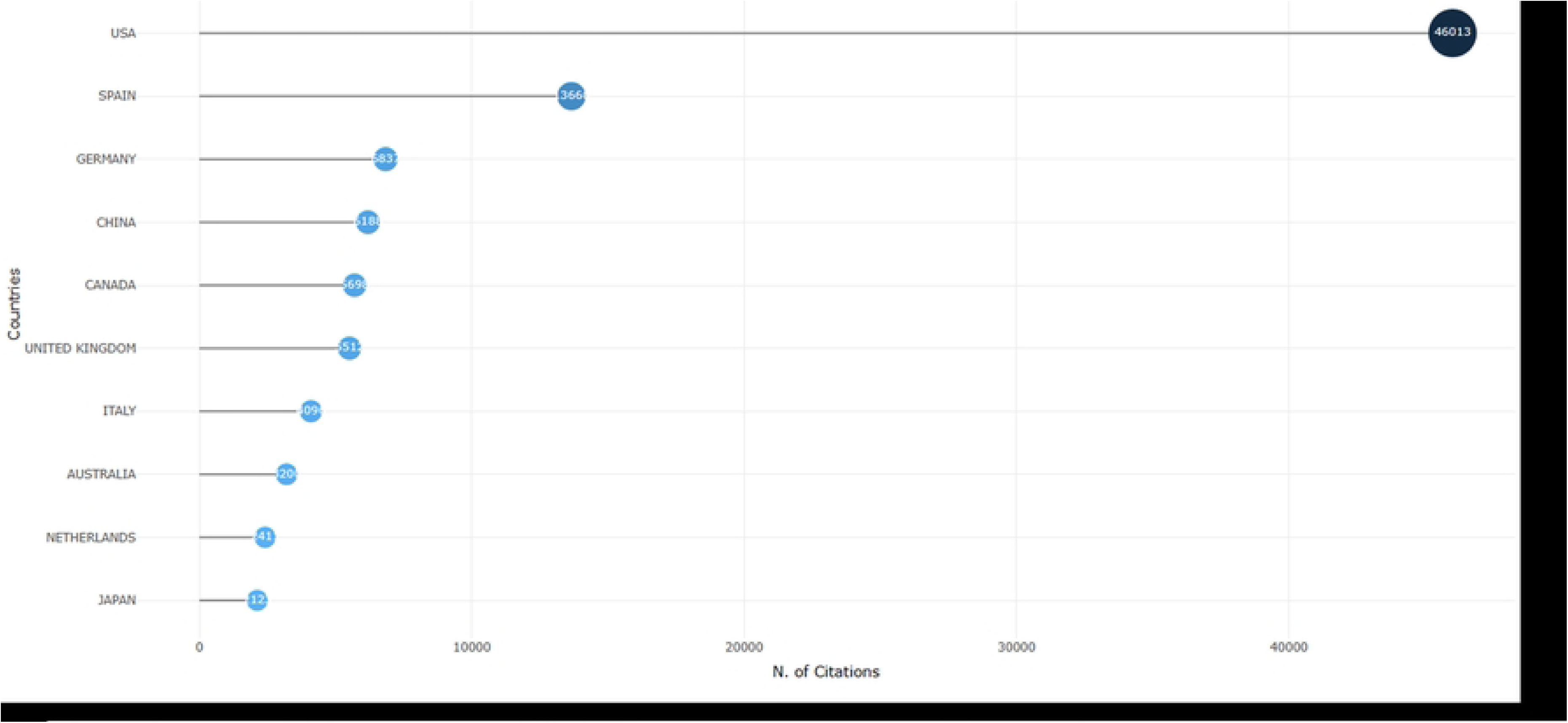
Quality of literature published in different countries/regions

### 2.2 A multidimensional analysis of the epigenetic age-related literature

#### 2.2.1 Research areas of the epigenetic age

As shown in Fig 4, in the research areas of the epigenetic age, the top 10 research areas in terms of the number of articles published within the WoS database are, with the five main research areas being cell biology (475), genetics (456), molecular biology (405), oncopathology (315) and integrative sciences (267).

**Fig 4.**
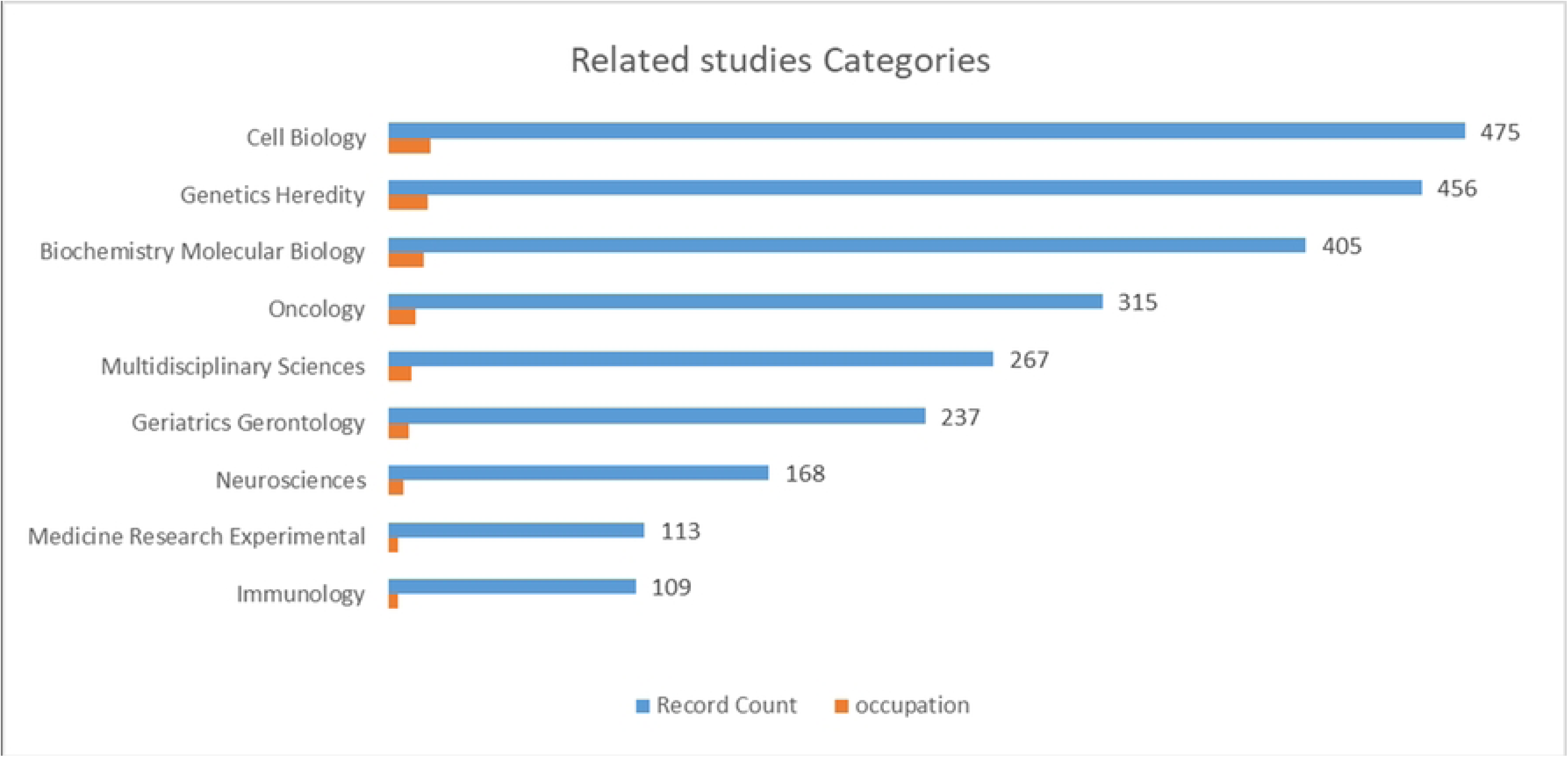
Epigenetic Age Research Area

#### 2.2.2 Key Institutions for Researching the Epigenetic Age

Conducting scientific research in the field of medicine requires a good research platform. Scientists need the help of research organizations to conduct scientific research. Fig 5 shows that the top five institutions in terms of number of publications are UNIVERSITIES OF CALIFORNIA SYSTEM, UNIVERSITY OF CALIFORNIA L OS ANGELES, HARVARD UNIVERSITY, JOHNS HOPKINS UNIVERSITY, UNIVERSITY OF LONDON, all from the United States and the United Kingdom.

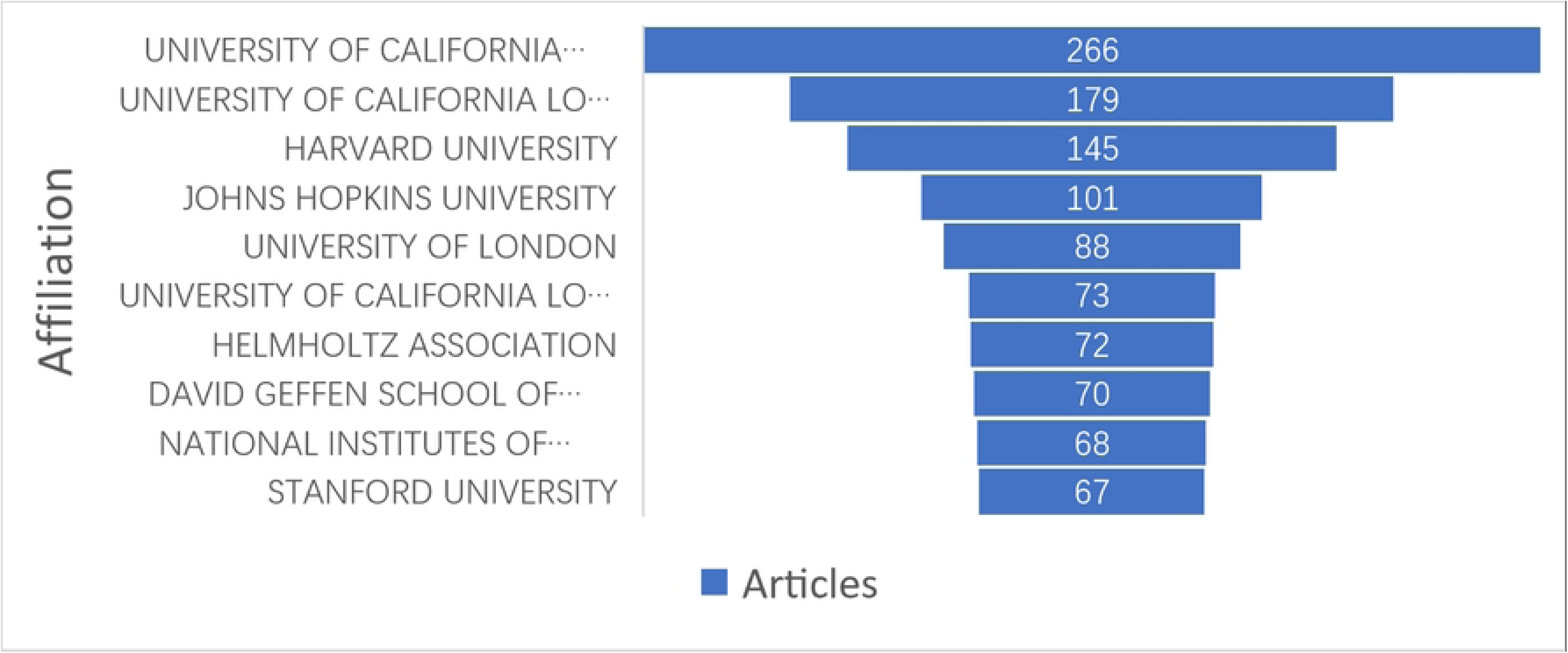

#### 2.2.3 Key authors studying epigenetic age

The Author Research Collaboration Network visualization map helps provide information about influential research teams and potential collaborators, thereby informing research groups about conducting collaborations. Fig 6 lists the top 10 authors with the highest number of publications in epigenetic age-related research areas. They are more active in this research area, with HORVATH S being the lead author with 41 publications. Among these authors, the largest number of authors are from the United States, indicating that the United States has paid more attention to epigenetic aging research and has a certain research base;

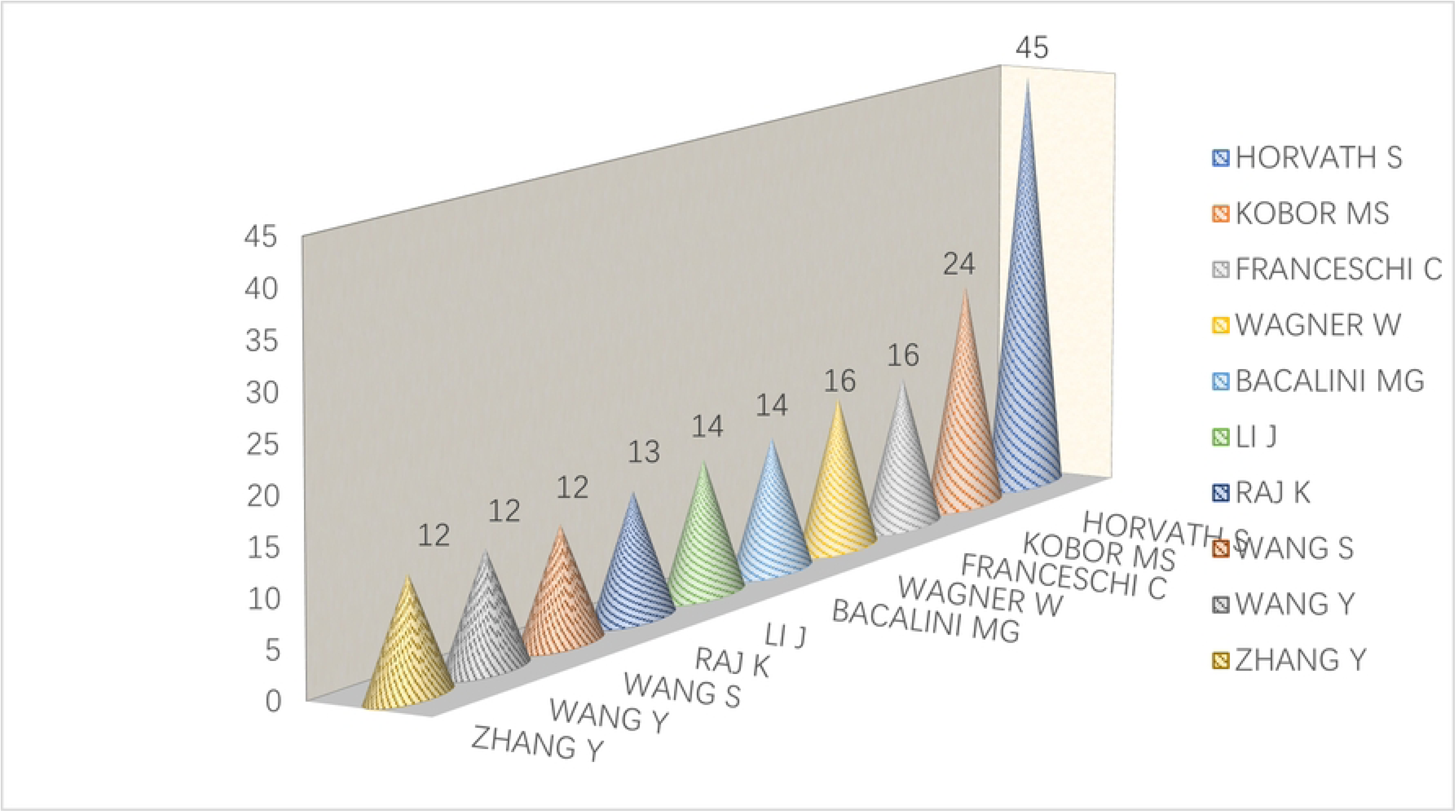

The topic of her research is mainly concerned with the study of genetic epigenetic age methylation.

#### 2.2.4 Major funds that finance epigenetic aging research

Fig 7 lists the top ten sponsors of epigenetic age-related research publications. The U.S. Department of Health and Human Services ranked first with 1,265 publications on epigenetic age; the National Institutes of Health ranked second with 1,260 publications; and the National Cancer Institute of the National Institutes of Health ranked third with 180 publications.

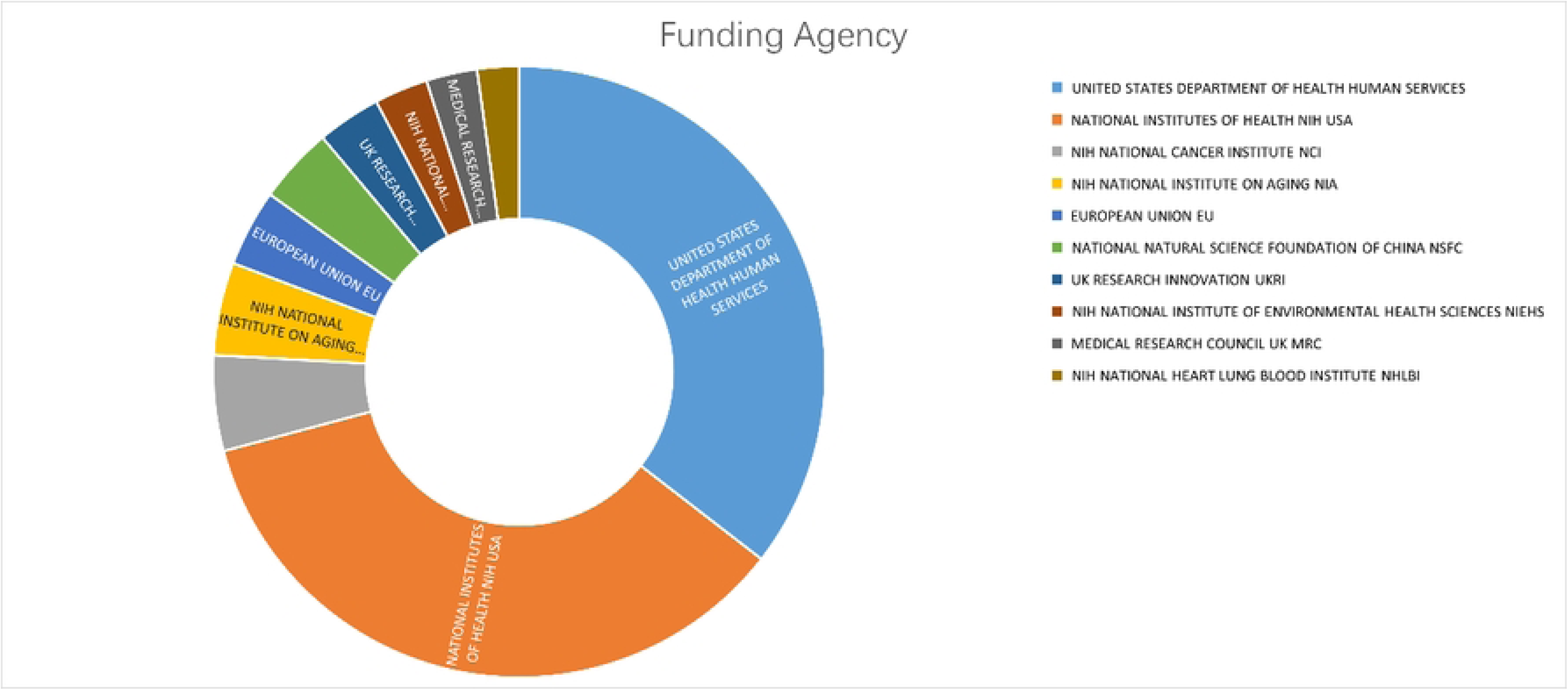

#### 2.2.5 Journals and periodicals publishing epigenetic age

SCI core journals are globally influential journals with a high number of articles and high circulation. Analyzing the distribution of core journals in the field of research related to epigenetic age can guide article submission. Fig 8 shows that the top three journals in terms of the number of articles published are PLOS ONE, CLINICAL EPIGENETICS and INTERNATIONAL JOURNAL OF MOLECULAR SCIENCES, which published a total of 252 articles (42%), indicating that these journals are the most important journals for articles in this area. By analyzing the most cited journals, it is possible to identify journals in the field that have a high impact and are hotly pursued for tracking the latest literature information and submitting articles. NATURE is the most frequently cited journal, and the relevant literature it publishes corresponds to the basics of research in the field, making it an important reference journal in the field.

**Fig 8.**
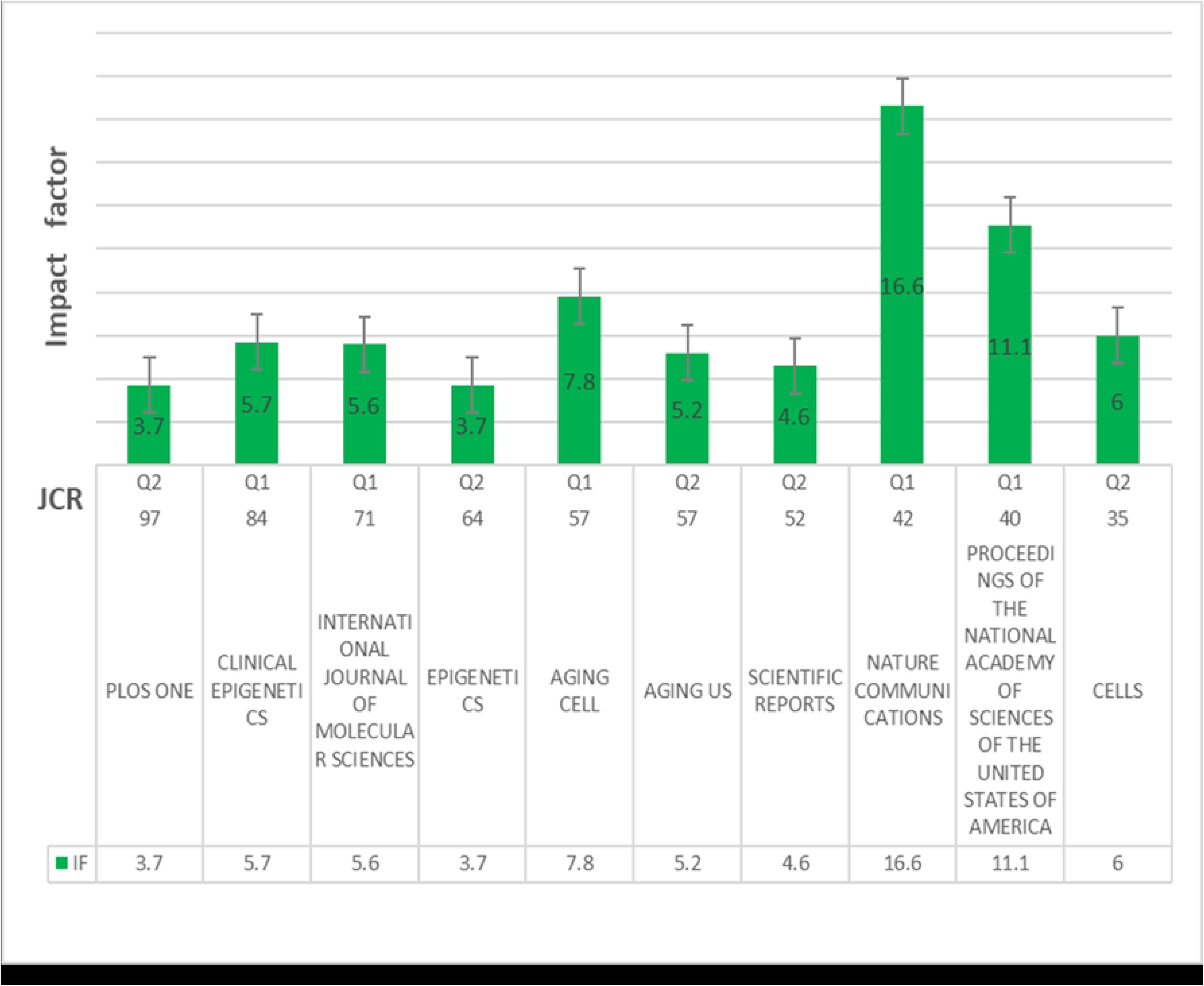
Journals and magazines that publish epigenetic age

### 2.3 Visual analysis of the literature on genetic epigenetic age

Co-occurrence analysis: In bibliometrics, co-occurrence refers to the co-occurrence of keywords, authors, institutions, etc. in multiple documents; Co-occurrence analysis is a quantitative study of simultaneously occurring phenomena to reveal the content association of the information and knowledge implied by the characteristic elements, which can help find out the direction of current research and the main focuses of research, and has a certain guiding effect on the Development of scientific research.

#### 2.3.1 Co-authors and co-author countries

Co-author analysis provides an understanding of the researchers conducting collaborative research in the field. Using VOSviewer software with the inclusion criterion of more than 5 co-authors, Fig 9 shows that a total of 87 authors were included in this analysis. Fig 9A shows the co-authors’ countries and the co-authors’ publication status. The three countries with the most co-authors are the United States with 661 articles (SCP 481 articles, MCP 180 articles). China ranks second behind the United States with 232 articles (SCP 169 articles, MCP 63 articles) and Germany with 77 articles (SCP 77 articles, MCP 47 articles), with SCP for single-country publication and MCP for multi-country co-publication is shown in Fig 9A. publication, and MCP stands for multi-country co-publication. Fig 9B shows the top five scholars with a high number of co-authored publications in the area of epigenetic age-related areas, with the top five researchers according to the authors in Fi g 9A Strength of overall connection between scholars include Horvath Steve, Franceschi Claudio, Bacalini Maria Giulia, Garagnani Paolo, KoborMichaels. The different colors in Fig 9B of the circles indicate different clusters. The number of circles indicates the number of co-authors, while the size of the circles indicates the size of the authors’ contributions, and the thickness of the lines between the co-authors indicates how closely related they are to each other.

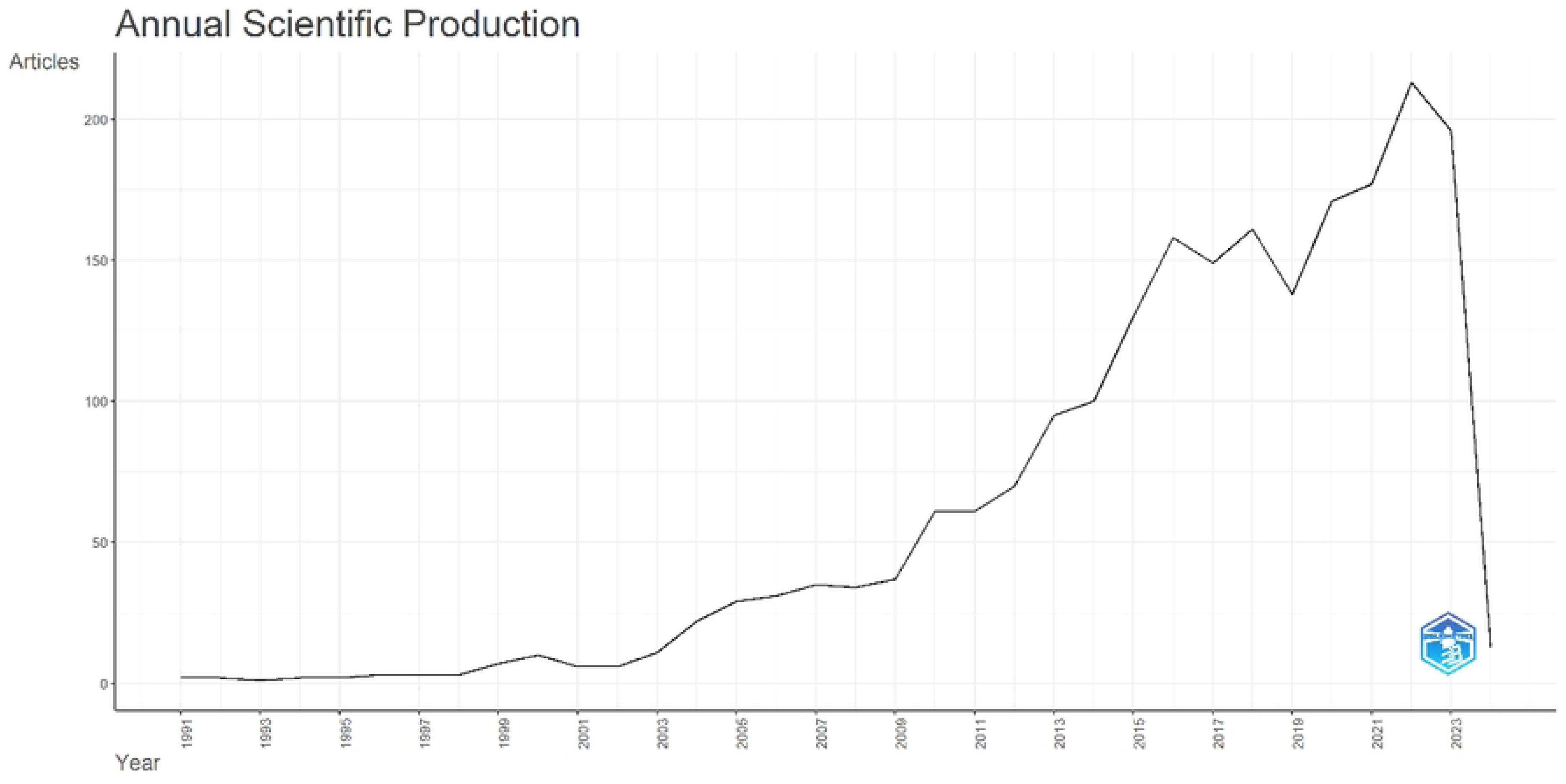

#### 2.3.2 Co-author of literature related to epigenetic age

In the CiteSpace software, a total of 2,800 institutions that published relevant literature were included in the co-author analysis, and it was determined that the institutions should publish at least 10 pieces of literature, and a total of 99 institutions met the conditions through the compilation, as shown in Fig.10 shown. According to the co-author institution statistics of the co-author analysis in the CiteSpace software, the five institutions with the highest co-author intensity of the published literature in this area were, in order of priority, as follows, UNIVERSITY OF CALIFORNIA SYSTEM, UNIVERSITY OF CALIFORNIA L OS ANGELES, HARVARD UNIVERSITY, JOHNS HOPKINS UNIVERSITY, UNIVERSITY OF LONDON. Different color circles indicate different clusters and circles. The number of co-authored organizations is indicated by the number of rows, while the size of the circle indicates the size of the contribution made by the organization and the number of rows between co-authored organizations indicates the degree of closeness between them.

**Fig 10.**
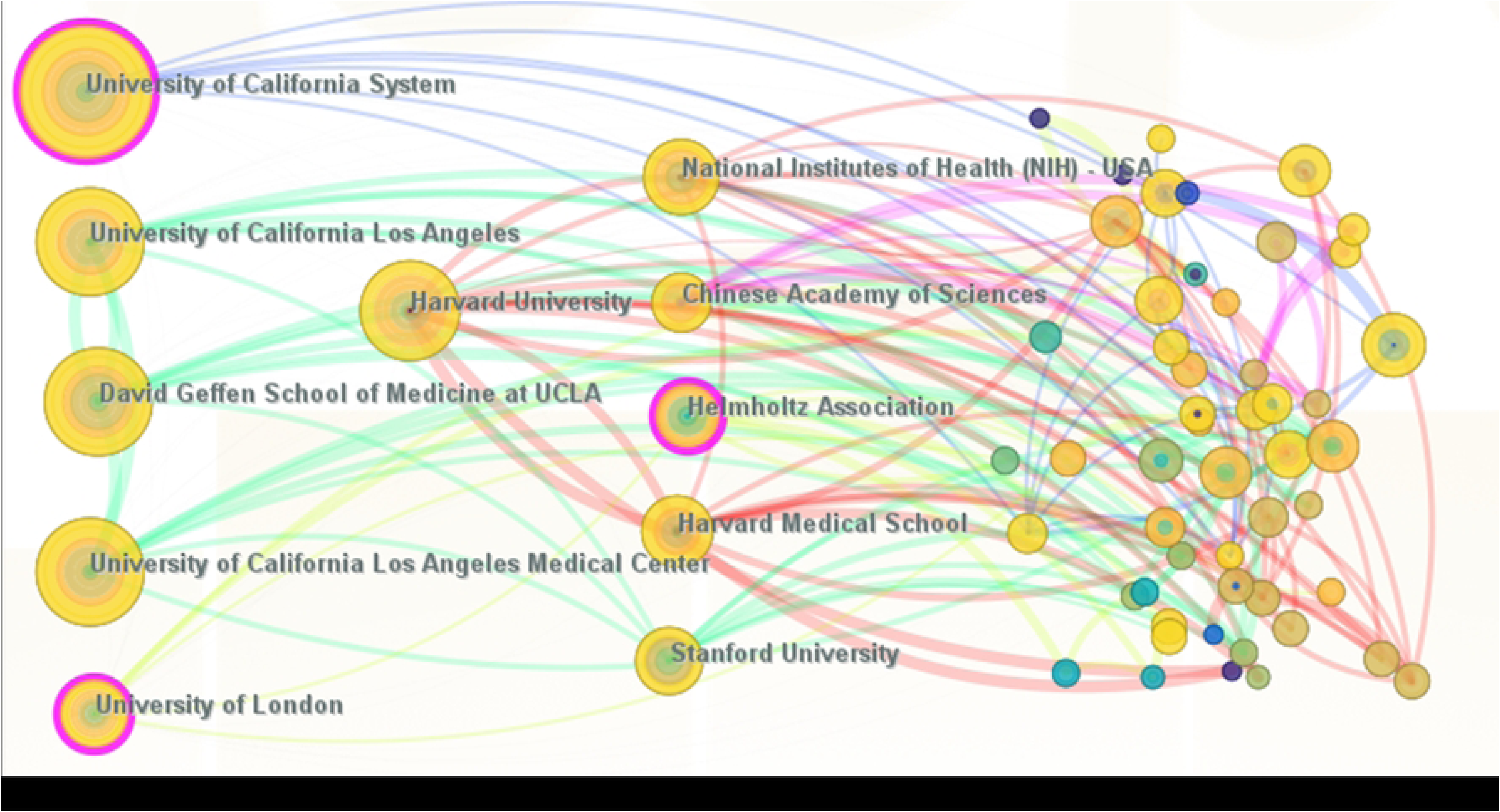
Co-authorship analysis chart of epigenetic age-related literature institutions

#### 2.3.3 Research directions related to epigenetic age

Using the co-occurrence analysis function of Citespace software, a total of 9229 keywords were used, and 829 keywords with a co-occurrence number of 5 or more were included as the inclusion criterion (as shown in Fig. 11).The current research directions of epigenetic age can can be divided into seven categories: the influence of human tis sues, the analysis of the genetic clock, the relationship between age-related diseases, precise genetic changes, accelerated cellular senescence, local cell transformation and genetic immune regulation. In order to further refine the analysis of the seven research directions, the five most frequently occurring keywords related to effects on human tissue cells, blood vessels, tissue, memory and birth; When analyzing the genetic clock, the key words genetic biomarkers, molecular aging, gene expression, and biological age appeared most frequently; Among ag e-related diseases, the keywords bone marrow, instability, hip fracture, and cognitive aging appeared most frequently; in the precision of genetic changes, The key words for age-related diseases are bone marrow, instability, hip fracture and cognitive aging. Aging; Among the precise genetic changes, diet, RNA methylation, oxidative stress and blood histones were more common. When it comes to accelerated cell aging, the key words that come up frequently are bone mar row, bone, osteogenesis, hematopoiesis, acetylation. In localized cell transformations, there is often a correlation between keywords: stromal cells, mesenchymal stem cells, histone acetylation, reactive oxygen species. In genetic immune regulation, binding site translocation, zinc regulation, mucus, oral, mitochondrial, dis mutase and *P53* are the most common.

**Fig 11.**
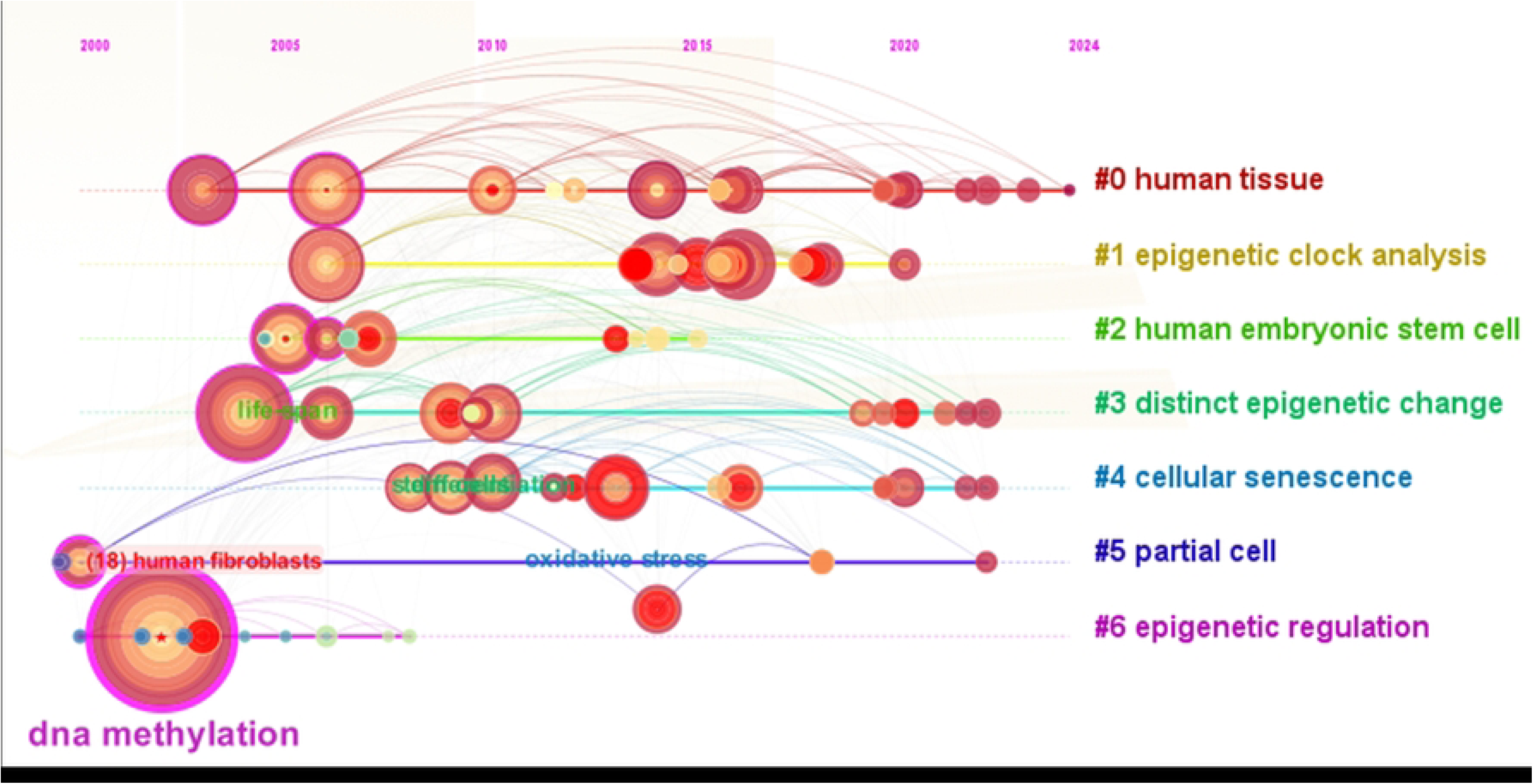
Diagram analysing the direction of epigenetic age-related research

#### 2.2.3 Trends and research hotspots of epigenetic aging research

The current research directions from 2000 to 2024 were extracted using the trend analysis function of Vos viewer software, and 8107 keywords were statistically analyzed, of which 168 keywords appeared at least 20 times, and the keywords that appeared more frequently in important ones time periods summarized. As shown in Fig. 12A, the research direction on genetic epigenetic age changes with the period, the color change scale in the lower right corner of the Fig is arranged from blue-light blue-green-yellow-orange-yellow, the blue part indicates that the keywords appeared earlier, which belongs to the previous research hotspot, and the yellow part represents the current research hotspot over time. The blue part of Fig 12A indicates that the research The research on epigenetic age has been focused since long on hypermethylation and promoter methylation. The yellow part of Fig 12A indicates that “oxidative stress”, “transduction pathway”, “mechanism study” and “activation response” are relatively new research directions. Statistical analysis using the online metrics platform bibliometrix in R software, as shown in Fig 12B, also concluded that the above four topics are currently recent research priorities. Keyword emergence analysis (12C) was carried out using Cite Space software. The results showed that “stem cells” were the most prominent topic in the epigenetic age literature over the past 20 years, and “calorie restriction” was the longest studied topic. In recent years, terms such as “lifespan,” “aging,” “profile,” “mechanism,” and “death” have emerged. In the future, these keywords may be a hot topic and a scientific breakthrough in studying the relationship between genetic factors, epigenetic age and human cellular aging.

**Fig 12A.**
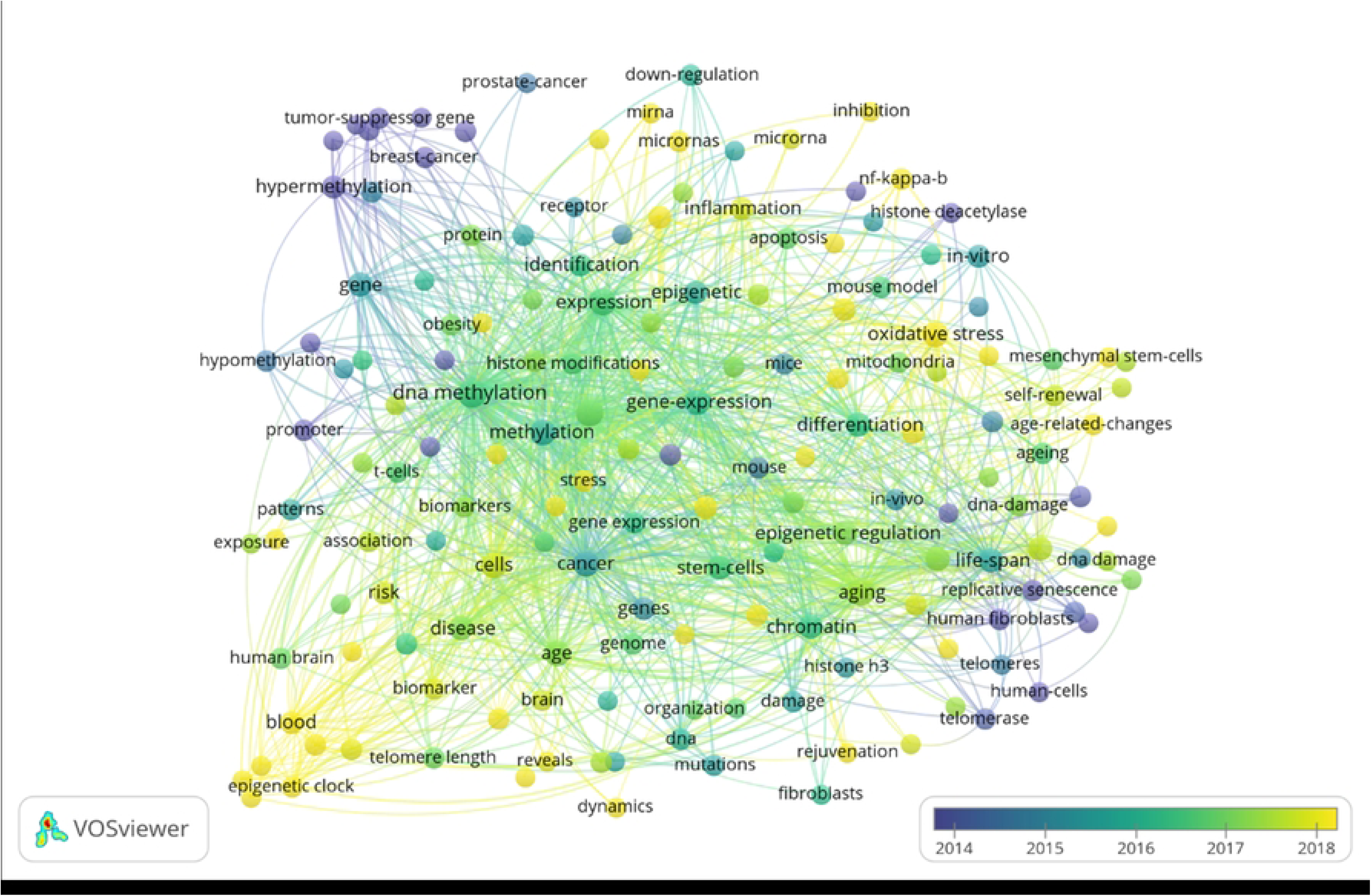
Trend analysis chart for epigenetic age studies

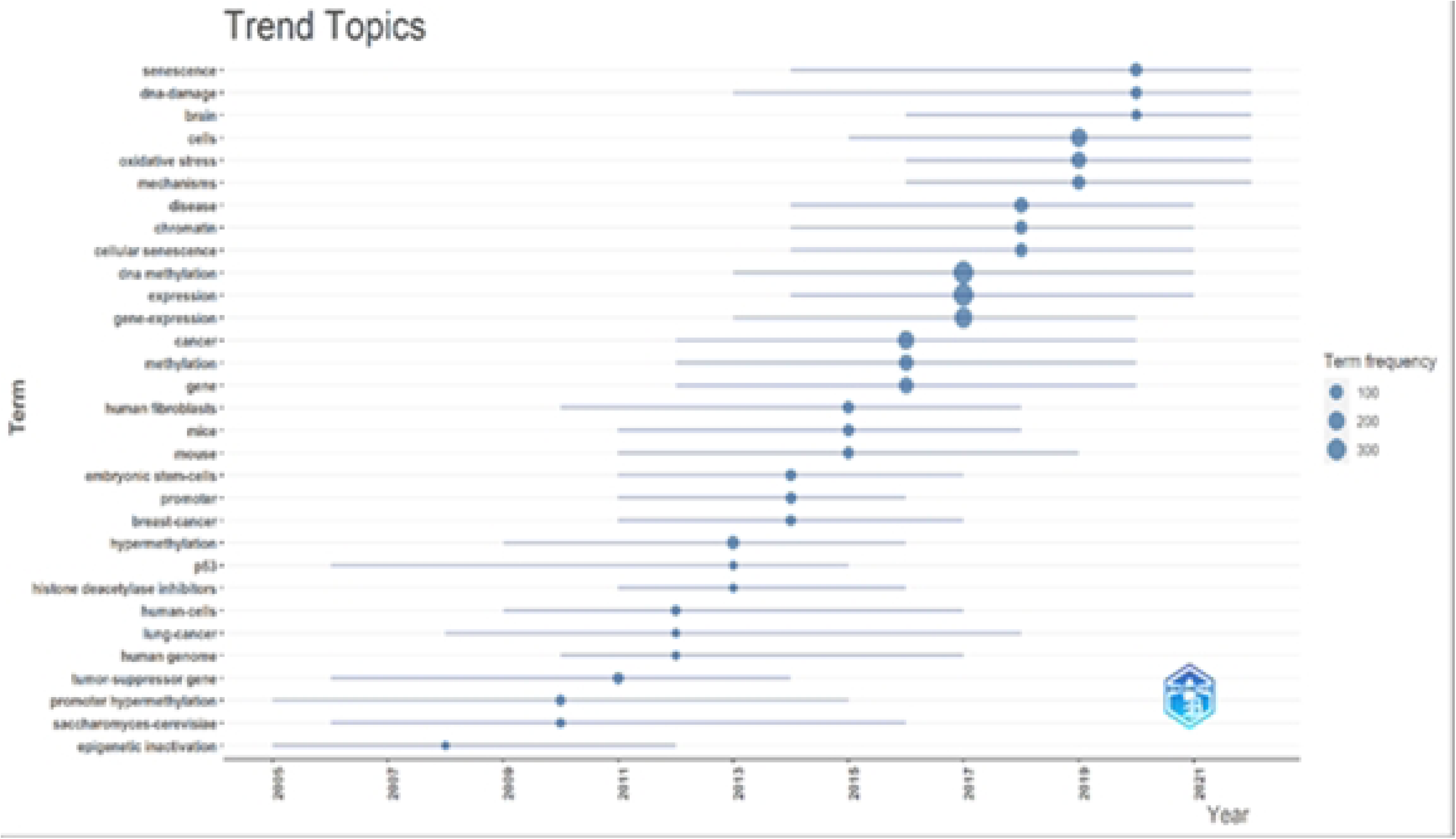

**Fig 12C.**
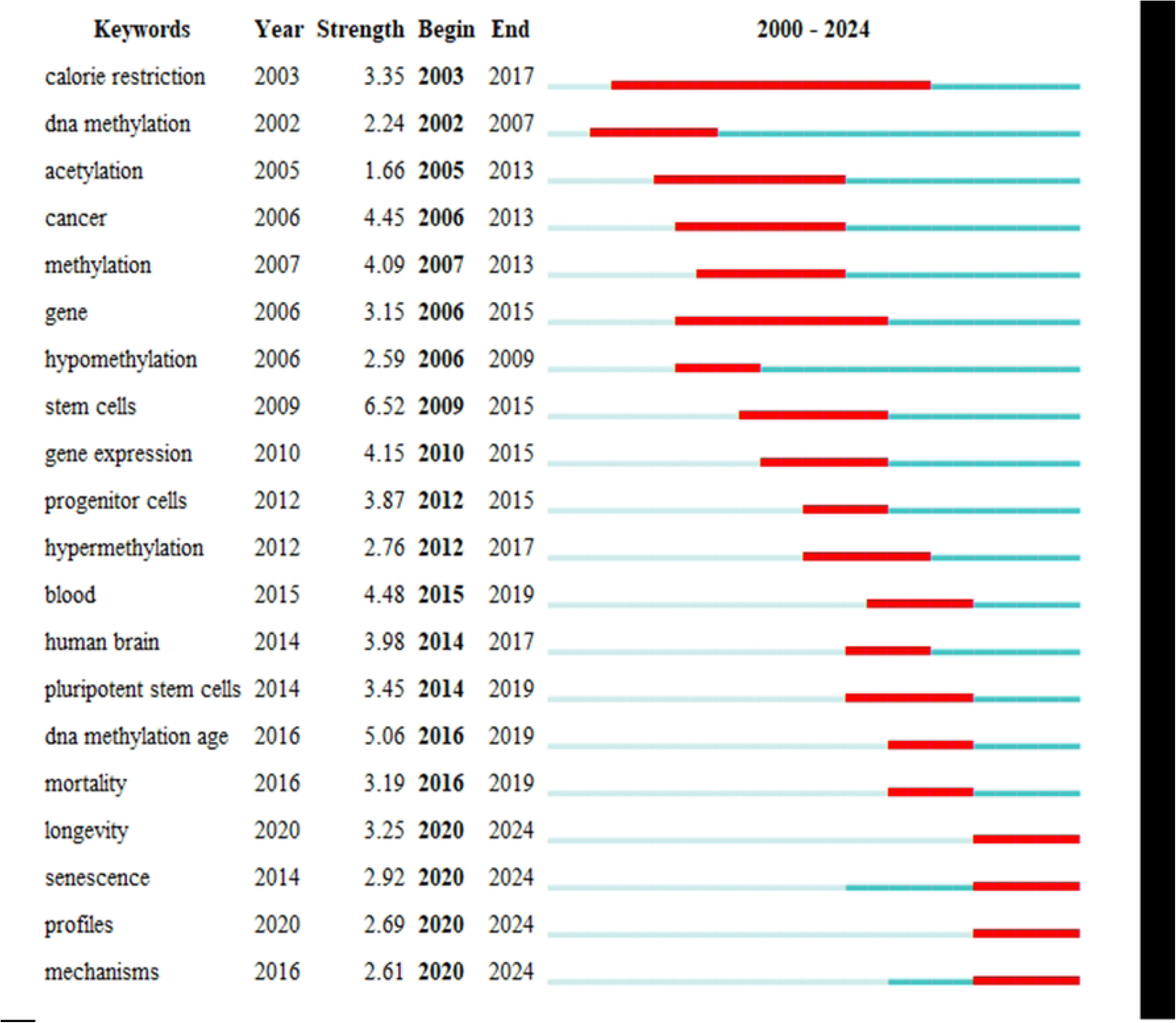
An analysis of future hotspots in epigenetic aging research Fig

## Discussion

In this study, we performed a bibliometric analysis of existing studies using the keywords “epigenetic age” and “cellular senescence”, with the aim of examining the relevant information in this area. According to the results, Most contributions to research on the relationship between epigenetic age and cellular senescence come from developed countries, and China, a developing country, is also on the list. Research in this area focuses on European, American and Asian countries. The field began in Europe and the United States, but there are also many developing countries that are actively engaged in this field at the same time. Keywords reveal the core topic and main content of an article, and the analysis of these keywords can adequately describe the research hotspots.

Cellular senescence is a condition in which cell proliferation is permanently inhibited and is a continuous irreversible process [11] with a variety of biological features, including increased senescence-associated β-galactosidase (SA-β-GAL) activity and a senescence-associated secretion phenotype [ 12] (SASP). Physiological aging is characterized by a gradual decline in physical or mental activity that leads to aging. Factors associated with aging include physiological age, genetic factors, lifestyle, diet, physical fitness, health status, social concerns, and cognitive functions [13]. Age is one of the major risk factors for man y age-related diseases [14], including cardiovascular diseases, endocrine diseases, cancer, immunodeficiency, bone metabolism, mitochondrial dysfunction, neurodegenerative diseases and immune-related diseases [15]. For example, senescent cells contribute to physiological aging and age-related diseases by causing a low-grade inflammatory state (senescence-associated secretory phenotype - SASP)[16]; senescent cells can stimulate cancer development through pro-inflammatory cytokines[17] ; Aging vascular cells, including endothelial cells and smooth muscle cells, are involved in atherosclerosis [18] ; and senescent preadipocytes and adipocytes cause insulin resistance [19] . Therefore, regulation of cellular senescence has been recognized as a potential anti-aging or delayed aging strategy [20]. Although current medical technology has advanced and significantly improved the survival rate of patients, the irreversible nature of aging also endangers human health and ultimately leads to death [21].

For decades, genes have been thought to determine the various proteins required in life and determine the phenotype of living organisms[22]. However, there are individual differences in the health of cells at different ages[23], and the trend is increasing. A number of research papers on cellular senescence are based on telomeres protecting the ends of chromosomes and on the shortening of telomere length with increasing age Age and the effects of factors that ultimately lead to the cessation of cell division[24]. Complementary to telomeres is epigenetic age, with telomere length and epigenetic patterns showing a correlation with the senescence characteristics of human cells [25]. Nine key factors have been demonstrated to contribute to aging, namely genomic instability, telomere depletion, loss of proteostasis, altered nutrient sensing, cellular senescence, mitochondrial dysfunction, stem cell depletion, altered intracellular signaling, and immune dysregulation and epigenetic changes [26]. .Therefore, in recent years, epigenetics has gradually become a research focus, playing a regulatory role through DNA methylation [27], Histone modification [28] (through acetylation/ubiquitination/methylation, etc.) and non-coding RNA mechanisms [29], particularly exploring the interactions of epigenetic age with cellular senescence in human cells, an d these studies may have the potential to diagnose diseases at an early stage and to improve health have significant impact [30].

Through collinear analysis of keywords, we can see that “DNA methylation” is a keyword that cannot be ignored. The most common form of DNA methylation is the addition of methyl groups to the 5′-cytosine of C-G dinucleotides, so-called CpGs [31]. These nucleotide pairs have low densities in the genome, and regions with relatively high CpG densities are known as CpG islands [32], identified as regions with >200 bp, >50% G + C content and 0.6 observed/expected CpGs ratios [33]. DNA methylation has received great attention in the study of genetic epigenetic age and cellular senescence, mainly due to the role it plays in the study of genetic epigenetic age and cellular senescence through various modifications of oxidative stress, DNA damage, Telomere changes and SASP development in cellular senescence [34]. For example, the cell cycle is regulated by changes in DNA methylation levels during telomere shortening [35]., which can affect telomere length and integrity to regulate cellular senescence, leading to DNA damage and promoting p53/p21 expression [36].During SASP secretion, DNA methylation affects the inflammatory process mainly by regulating the expression of pro- and anti-inflammatory cytokines [37], which in turn affect the cell cycle by regulating p53/p21 and p16 expression, thereby influencing the progression of cellular senescence [38].During DNA damage, methylation modifications regulate cellular senescence by influencing DNA polymerase recruitment at damage sites and regulating DNA break repair to regulate p53/p21 expression [39].During oxidative stress, methylation mainly acts on oxidative and antioxidant systems to regulate oxidative and antioxidant homeostasis in vivo [40].

The research hotspots of genetic epigenetic age and cellular senescence research show different trends of temporal distribution. The focus of cellular senescence research has shifted from the pathological level to the molecular level, and the role of genetic epigenetic age has begun to receive attention from different perspectives. Interdisciplinarity and holistic development have characterized the research process. In addition to focusing on how epigenetic age is involved in the progression of cellular senescence, others remain focused on the study of the mechanisms of senescence and on therapeutic targets.

## Research limitations

This study collected relevant literature on epigenetic age and cellular senescence from 2000 to 2024, and the relevant literature in the WOS core collection will be updated as more in-depth studies are conducted in later years, so the results will be retrieved from this Study may differ slightly from the actual number of literature. During the literature collection process, only the Web of Science core collection was used to conduct a systematic literature search, and literature published in other databases may be omitted, which may affect the accuracy of the results of the study. Due to the different steps involved in using visualization software and the fact that each software has its own visualization and calculation characteristics, some important details may be missed during analysis. The language of the literature was only used as the only search criterion, so some high-quality non-English literature data may be missed during the search process.

## Conclusion

In summary, the results of bibliometric and visualization analyzes revealed that publications on the connection between epigenetics and cellular senescence have increased year by year worldwide over the past two decades. Regulation of cellular senescence is considered a potential anti-aging strategy to prevent cellular senescence and influence the secretory phenotype through continuous optimization of existing research techniques, but also research related to the removal of already senescent cells and the epigenetic reprogramming of senescent cells. Therefore, there is a greater need to provide the latest evidence for anti-aging therapies through prospective, multicenter, high-quality clinical trials.

## Supporting information

**S1 Fig. Trends of Cellular Senescence and Epigenetic Age publications over the past 20 years.**

**S2A Fig. Distribution of the top 10 countries / territories in global publication rankings**

**S2B Fig. Number of publications in different countries or regions**

**S3 Fig. Distribution of the total number of citations in the top 10 countries / regions in the global publication rankings**

**S4 Fig. Epigenetic Age Research Area. The top 10 main research directions in this field**

**S5 Fig. Key Institutions for Researching the Epigenetic Age. Institutions with the world ’s top 10 publications in this field**

**S6 Fig. Key authors studying epigenetic age. Authors of the world ’s top 20 publications in this field**

**S7 Fig. Major funds for epigenetic age research. Funds that rank among the world ’s top 20 in the number of publications in this field**

**S8 Fig. Journals and magazines that publish epigenetic age**

**S9 Fig. Country analysis chart of co-authors of epigenetic age-related literature**

**S10 Fig. Co-authorship analysis chart of epigenetic age-related literature institutions**

**S11 Fig. Diagram analysing the direction of epigenetic age-related research**

**S12A Fig. Trend analysis chart for epigenetic age studies**

**S12B Fig. Hot trend analysis map for epigenetic age studies**

**S12C Fig. An analysis of future hotspots in epigenetic age research Fig**

## Acknowledgments

The authors wish to thank Hao Wang and Zhi-Guo Sun for helpful advice and discussions.

## Author Contributions

Conceptualization: Mureti Kahaer, Dilimulati Aikeremu, Hao Wang

Datacuration:Mureti Kahaer

Formal analysis: Dilimulati Akeem, Kuo Xu

Investigation: Zhi-Guo Sun

Methodology: Mureti Kahaer, Kuo Xu

Project administration: Hao Wang

Resources: Dilimulati Aikeremu

Software:Mureti, Zhi-Guo Sun

Supervision: Hao Wang,

Validation: Zhi-Guo Sun, Kuo Xu

Writing –original draft: Mureti Kahaer.

Writing –review&editing: Dilimulati Aikeremu, Zhi-Guo Sun

---

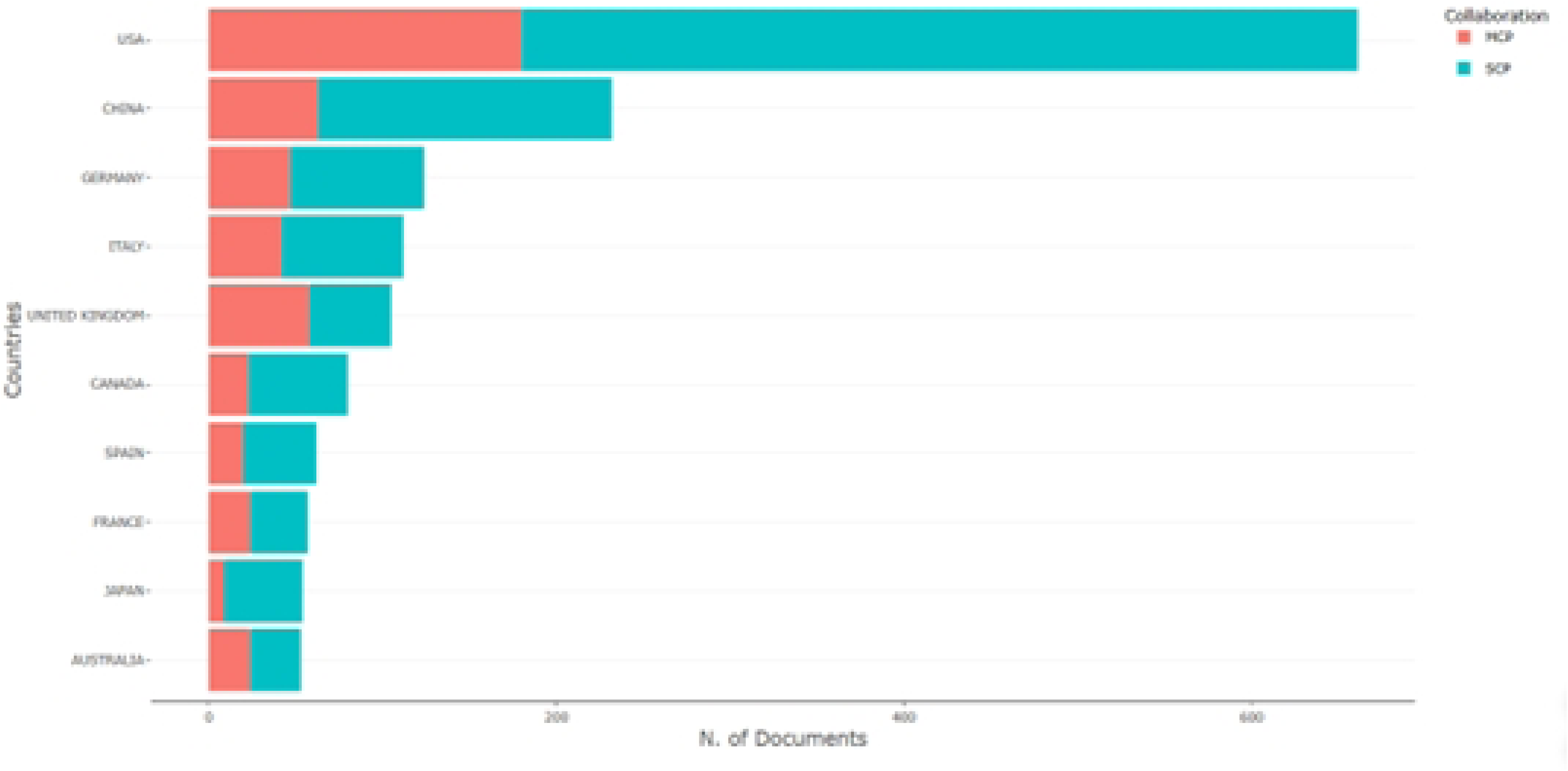

